# Single-cell RNA sequencing analysis reveals new insights for intersex gonad initiation and early sex differentiation of goats

**DOI:** 10.1101/2024.12.31.630861

**Authors:** Xinxin Cao, Yichen Zhang, Dongliang Xu, Mengmeng Du, Xiaokun Lin, Lu Leng, FM Perez Campo, Lihui Zhang, Mingzhi Sun, Xiaoxiao Gao, Jianning He, Qinan Zhao, Jianguang Wang, Jinshan Zhao, Hegang Li

## Abstract

Polled intersex syndrome in goats severely impacts productivity and economic benefits, serving as a potential model for human reproductive defects. However, its specific molecular and cellular regulatory mechanisms remain unclear. This study combined single-cell RNA-sequencing (scRNA-seq) with tissue morphology and immunofluorescence detection to investigate the cellular heterogeneity, pseudo-time differentiation trajectory, differentially expressed genes, and regulatory networks of female, intersex, and male gonads during the 55-65 days embryonic stage of goats. The results indicated significant heterogeneity and cell fate transition in the somatic and germ cells of female, intersex, and male gonads at different developmental stages. Stromal cells may initiate female and intersexual gonad development, and epithelial cells may be the source of gonad cells in male fetuses. The abnormal expression of *FOXL2*, *SOX9*, *AMH*, *DMRT1*, *RSPO1*, *INSL3*, *TCF21* and *CSF1R* may cause female-to-male sex reversal. The new candidate genes like *PHLDB2*, *ZNRF3* and *TEAD1* may contribute to the gender differentiation. Additionally, *TREM2^+^* macrophages analogous to those in humans were identified in male gonads, while genes such as *CSF1R* and *TYROBP* potentially regulate macrophage immune privilege mechanisms. Early gonadal development in goats may also be regulated by circadian rhythm mechanisms similar to humans, with genes such as *NR1D1*, *LGR4*, and *ATG5* involved in granulosa cell development, while genes such as *ATG5*, *NRIP1*, and *FBXW11* may affect intersex trait development. In general, this study provided an important reference for exploring molecular and cellular mechanisms of intersex initiation during early embryonic development in goats and also for modelling human reproductive defects.

**Author summary:** In this study, we investigated the cell fate trajectory and key gene expression profile of goat fetus gonads using scRNA-seq. The analysis revealed cellular heterogeneity among normal male, normal female, and intersex gonadal somatic and germ cells, alongside their dynamic fate transitions across various developmental states. Pseudo-time trajectory analysis suggested that stromal cells might act as the initiation signal for female and intersexual gonad development, while epithelial cells could serve as the primary source of gonadal cells in male fetal goats. GO functional enrichment and KEGG analyses confirmed that abnormal expression of *FOXL2*, *SOX9*, *AMH*, *DMRT1* and *RSPO1* was linked to the emergence of intersexual gonads. Newly identified candidate genes including *LRP2*, *PITX2*, *ARX*, *LHX9*, *PHLDB2*, *ZNRF3* and *TEAD1* may contribute to sex-specific differences, while *INSL3* and *TCF21* could be linked to intersexual gonads. We identified *TREM2^+^* macrophages analogous to those in humans, which could regulate immune privilege mechanisms. Circadian rhythmic changes were observed during early goat gonadal development.

## Introduction

In mammals, intersex or sex-reversal phenomena can occur during sexual differentiation [1]. Intersex refers to atypical sexual differentiation or dysplasia that results in the absence of normal female gonadal function or a reversal in morphology. Intersex individual often suffers from congenital reproductive disorders, which are generally associated with slower growth rates and higher mortality [2,3]. The phenomena seriously affect the expansion and production efficiency of the goat population severely impacting the development of the goat industry [4]. The intersex trait in goats is linked to the polled character, referred to as Polled Intersex Syndrome (PIS) [5]. The PIS mutation region has been precisely mapped at chromosome 1q43, spanning 10,159 bp, and contains a reverse insertion of approximately 480 kb of repetitive sequences [3,6,7]. The transcription factor *FOXL2*, which plays a critical role in ovarian differentiation and development, is located approximately 200 kb downstream of the PIS sequence and is the only protein-coding gene regulated by the PIS segment [8,9]. Moreover, Blepharophimosis-ptosis-epicanthus inversus syndrome (BPES) in humans, which is associated with female infertility, is caused by pathogenic variants of *FOXL2* [10–13]. The translocation breakpoints in BPES patients are located in the chromosome region 3q23 [14], which shown significant homology with the locus on chromosome 1q43 containing the PIS region in goats [6]. This suggests that the molecular mechanisms underlying intersex traits in goats and human reproductive disorders share similarities, making intersex goats a potentially valuable model for studying human infertility. Consequently, understanding the mechanism of intersex formation in goats holds great significance for advancing animal husbandry and improving human reproductive health.

The formation of gonads is a crucial step in the sex determination process of mammals [15]. The genes *SF1*, *WT1* and *LHX9* are essential for the transformation of the genital ridge into a bipotential gonad in mice [16]. The development of bipotential gonads into either male or female gonads depends on the disruption of the balance between antagonistic and repressive signaling pathways within [17]. The somatic chromosomal composition determines whether the gonadal ridges develop into ovaries or testes [18,19]. *SRY* triggers the expression of *SOX9* and induces male-specific differentiation of mouse fibroblasts into supporting cells, facilitated by *GATA4*, *WT1*, *DMRT1* and *SF1* [20–23]. In the absence of *SRY*, the *WNT4*/*RSPO1*/*CTNNB1*/*FOXL2* pathways are activated, leading to the emergence of granulosa cells [18,24,25]. During XX sex reversal, *SOX9*, with the assistance of *DMRT1*, binds to accessible genomic regions [26]. *DMRT1,* which is expressed in goat supporting cells, plays multiple roles, including male maintenance, apoptosis regulation and male determination [27–32]. Knocking out *DMRT1* in male mice leads to extensive conversion of supporting cells into granulosa cells [33,34]. However, the molecular mechanism underlying intersex malformation in goats remains unclear. Although *FOXL2* has been identified as the primary gene responsible for female sex determination in goats [35], the specific molecular mechanisms driving the formation of intersex reproductive defects are not well understood, particularly in terms of the cell types involved during the embryonic stage. Further research is needed to elucidate the precise signaling pathways through which the *FOXL2* gene governs the process of female sex determination.

Primordial germ cells (PGCs), the direct precursors to oocytes and spermatogonium, typically migrate from the periphery of the blastocyst to the developing testes and ovaries through a combination of active and passive movement [36,37]. In mice, PGCs originate from the proximal allantois and migrate through the hindgut endoderm to reach the gonadal ridges [38]. PGCs exhibit sex-dimorphic expression of female-specific genes before the transition to female development, with germ cells arresting at Pre-meiosis I [39]. Retinoic acid (RA) induces mouse PGCs to enter meiosis. Prior to meiosis, human oocytes upregulate the *ZIC1*, while *DMRTC2*, *ZNF711*, and *DMRTB1* are activated after entering meiosis [40].

In male gonads, germ cell mitosis ceases after sex determination [38]. The Nodal/Cripto signaling pathway is rapidly activated [41], leading to the downregulation of pluripotency genes such as *NANOG*, *SOX2*, and *POU5F1* [42,43], while simultaneously initiating the expression of genes promoting male germ cell differentiation. *GATA4*, a primate-specific transcription factor, is upregulated in PGCs, and the transition of PGCs to spermatogonia involves the activation of *EGR4*, *KLF6*, and *KLF7* [40]. *SOX4* deficiency increases the number of testicular cords and promotes regulation of male germ cell differentiation in mice [44]. However, the heterogeneity of germ cells and related pathogenic genes in the early stage of intersex goat gonadal formation is still unclear.

Intersex traits in goats have been shown to manifest during the early stages of embryonic development [35,45]. However, the cellular heterogeneity of intersex gonads in goat embryos remains unclear, and this information is crucial to understanding the mechanisms underlying the formation of intersex traits. This study employed single-cell RNA-sequencing (scRNA-seq) on the early embryonic female, male, and intersex gonadal tissues of goats, aiming to explore the molecular and cellular regulatory mechanisms of goat intersex formation through comparative analyses. This research not only provides insights for correcting intersex traits, but also offers candidate disease models for studying human infertility.

## Results

### ScRNA-seq identified early germ cells and gonadal somatic cells in goats

To investigate the regulatory mechanisms underlying the differentiation of germ cells and gonadal somatic cells in early goat embryos, the transcriptome expression profiles of embryonic gonads were analyzed using single cell RNA-sequencing (sc-RNA seq) (Fig 1A). A total of 114,226 cells were estimated, and after stringent quality control, 97,214 high-quality cells were retained. The gonadal cells were categorized into two primary populations: germ cells and somatic cells. This study identified 4,393 germ cells (1,200 from female gonads, 998 from male gonads, and 2,195 were from intersexual gonads) and 92,821 somatic cells (27,481 from female gonads, 35,326 from male gonads, and 30,014 from intersexual gonads) (S1 Table).

**Fig 1.**
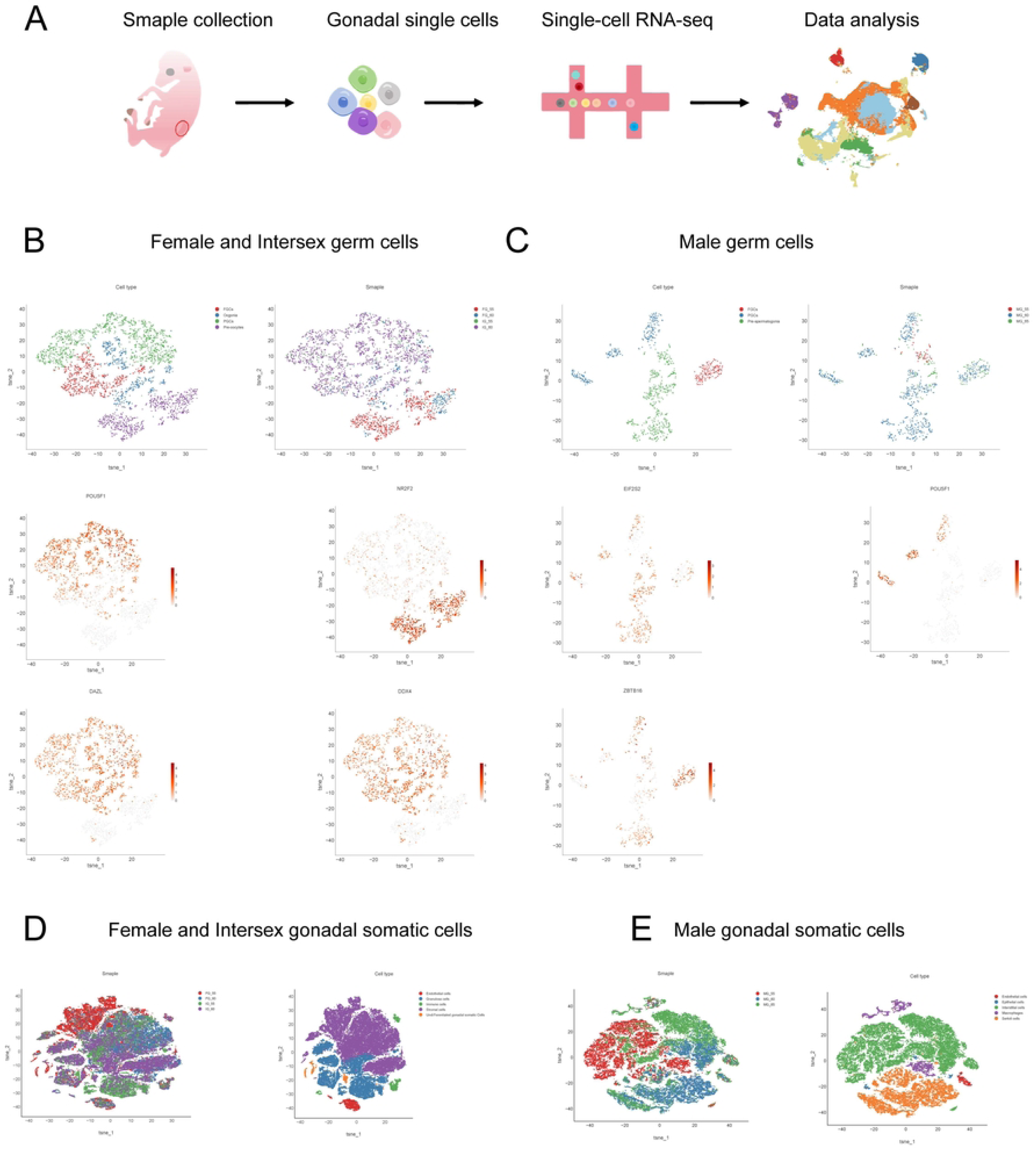
scRNA-seq delineated cellular heterogeneity during dairy goat gonad development. **A**, Overview of the experimental strategy. **B-C,** t-stochastic neighborhood embedding (t-SNE) showing 2D visualization of female, intersex (B) and male (C) germ cells labelled with developmental time and cell type. Cells from different developmental time points and cell types were color coded. Cells were color coded based on normalized gene activity. **D-E,** t-SNE showing 2D visualization of female, intersex (D) and male (E) gonad somatic cells labelled with developmental time and cell type. Cells from different developmental time points and cell types were color coded.

t-SNE clustering analysis was performed separately for gonadal somatic cells and germ cells. For intersex and female gonads, 21 and 8 clusters were respectively obtained from somatic and germ cells. For male gonads, 22 and 8 clusters were respectively obtained from somatic and germ cells (S1A Fig). Germ cells from female and intersex gonads were categorized into four subsets: primordial germ cells (PGCs), fetal germ cells (FGCs), oogonia, and pre-oocytes (Fig 1B). High *POU5F1* expression was observed in PGCs. FGC clusters (1 and 8) exhibited high expression of the marker gene *DDX4*. The marker gene *DAZL* showed elevated expression in oogonia clusters (5 and 6), while the *NR2F2* expression was prominent in pre-oocyte (2 and 4) (Fig 1B). Male germ cells were classified into three subsets: PGCs, FGCs, and pre-spermatogonia (Fig 1C). High *POU5F1* expression was observed in PGCs, while *ZBTB16* and *EIF2S2* were strongly expressed in FGCs and pre-spermatogonia, respectively (Fig 1C). Notably, the proportion of pre-oocytes in intersex gonads was higher than in normal female gonads, while the proportion of pre-spermatogonia in male decreased with age (S1B and S1C Fig).

Female and intersex gonadal somatic cells were classified into five cell types (Fig 1D): stromal cells (*TCF21*, *PDGFRB*), granulosa cells (*AMH*, *SERPINE2*), endothelial cells (*PECAM1*), immune cells (*AIF1*, *PTPRC*), and undifferentiated gonadal cells (*CITED2*) (S1D Fig). Stromal cells, (clusters 0, 1, 5, 6, 8, 12, and 17) exhibited high expression of *TCF21* and *PDGFRB*. Granulosa cells (Clusters 2, 3, 4, 7, 9, 10, 15, and 16) expressed *AMH* and *SERPINE2*. Endothelial cells (clusters 11 and 21) showed high *PECAM1* expression, while immune cells (clusters 13, 18, and 20) had high *AIF1* and *PTPRC* expression. Undifferentiated gonadal cells (clusters 14 and 19) displayed elevated *CITED2* expression.

Male gonadal somatic cells were also classified into five cell types (Fig 1E): epithelial cells (*KRT19*), interstitial cells (*PDGFRB*, *TCF21*), Sertoli cells (*AMH*), macrophages (*ACTC1*, *PTPRC*), and endothelial cells (*PECAM1*) (S1D Fig). Epithelial marker gene *KDR* was highly expressed in clusters 18 and 21. The endothelial cell marker gene *KDR* exhibited high expression in cluster 15. Sertoli cells expressed high levels of *AMH* in clusters 1, 5, 7, 8, 17, and 19. Stromal cells (clusters 0, 2, 3, 4, 6, 9, 10, 12, 14, 20, and 22) exhibited high expression of *PDGFRB* and *TCF21*. Macrophage marker genes *ACTC1* and *PTPRC* were strongly expressed in clusters 11, 13, and 16. Marker gene expression varied between females and males, indicating that differences in cell states during gonadal development are regulated by distinct genetic mechanisms.

### Dynamics gene expression and transcriptional regulatory network in germ cells of different ages

The enrichment of gonadal DEGs and their associated biological processes at various ages and across sexes were further compared (S2 Table). At 55 days post coitum (dpc), terms related to stem cell differentiation, sex differentiation and cell cycle processes were significantly enriched in both female and male embryos. However, intersex embryos showed enrichment primary for stem cell development (Fig 2A and 2B). At 60 dpc, females displayed enrichment for terms related to sex differentiation and meiosis, males were enriched for mitochondrial translation, and intersex embryos showed enrichment for the regulation of cell cycle processes and the mitotic cell cycle (Fig 2A and 2B). At 65 dpc, male embryos exhibited enrichment in stem cell differentiation, sister chromatid cohesion, and sex differentiation processes (Fig 2B).

**Fig 2.**
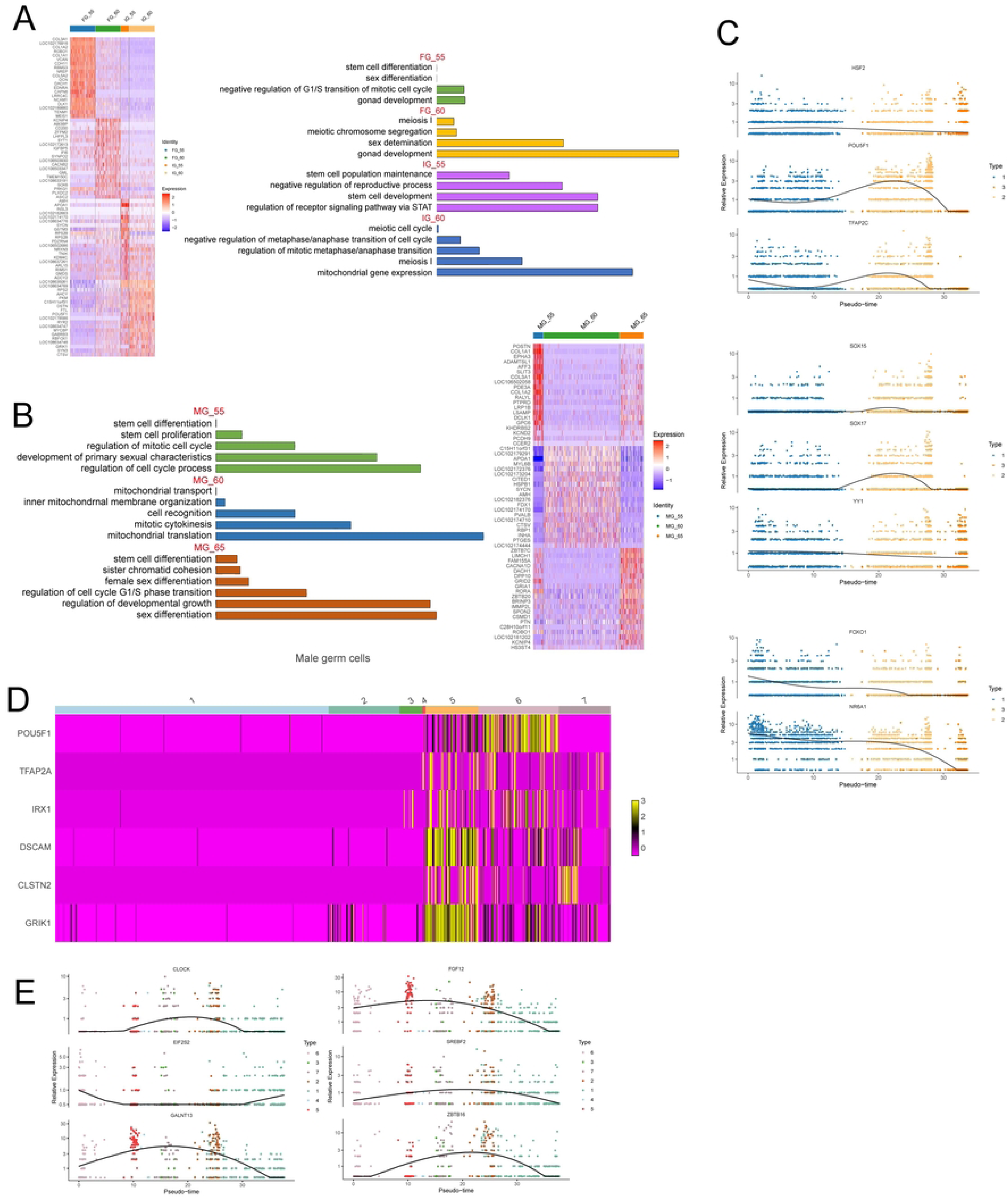
Dynamics gene expression and transcriptional regulatory network in germ cells of different ages. **A-B**, Heat map of DEGs expression in female, intersex, and male gonadal germ cells at different time points (left). Each grid represented the expression level of the smaple’s gene, ranging from red to blue, with decreasing expression levels. Enriched GO function (right). **C,** Visualization of female and intersex gonadal germ cells’ characteristic genes at different stages along pseudo-time. **D,** DEGs expression of male germ cells pattern heat map. **E,** Visualization of male germ cells’ characteristic genes at different stages along pseudo-time.

Notably, *ZEB2* was enriched in the cell surface receptor signaling pathway involved in cell-cell signaling in females, and in the regulation of the Wnt signaling pathway in MG_65. *POU5F1* expression was upregulated in intersexual goat germ cells, with no significant difference observed between female and male expression. In IG_60, *SOX17* was associated with cell division, while *SOX15* was involved in the regulation of cell cycle processes.

To further investigate the role of germ cell-related genes during development, the pseudo-time trajectories were reconstructed using transcriptomic data (S2A Fig). In female and intersex germ cells, genes such as *FOXO1* and *NR6A1* exhibited high expression during early developmental stages. In contrast, *POU5F1*, *TFAP2C*, and *HSF2* were expressed throughout development, with significant upregulation during later stages. SOX family genes, including *SOX17* and *SOX15*, displayed transient upregulation during mid-development. The differentiation-related gene *YY1* maintained a dynamic expression throughout gonadal development (Fig 2C).

Candidate genes associated with germ cell development were also identified. For instance, *ZEB2* may play a role in cell-cell signaling in females. *CLSTN2*, *DSCAM*, *FGF12*, *GALNT13*, *GRIK1*, *KCNJ13*, and *MED12L* were active throughout development but showed downregulation sat later stages, while *RPL24*, *RPL35A*, and *RPL37A* exhibited significant upregulation during later stages. *LRP1B*, *MBNL1*, *RBP1*, *ROBO1*, and *ZEB2* were highly expressed during later developmental stages (S2B Fig).

In male germ cells, *TFAP2A*, *IRX1*, *POU5F1*, *CLSTN2*, *DSCAM*, and *GRIK1* were specifically expressed in PGCs (Fig 2D), with high expression detected during early germ cells development (Fig S2C). *ZBTB16*, a marker gene for FGCs, along with *SREBF2*, *FGF12*, and *GALNT13*, showed specific expression across different developmental stages. *CLOCK* exhibited high expression during mid-development and was enriched in the intracellular receptor signaling pathway. *EIF2S2*, a marker gene for pre-spermatogonia, was upregulated during both early and later developmental stages (Fig 2E). *INSL3* was upregulated during FGCs and pre-spermatogonia development, while *NRXN3* showed upregulation during the development of PGCs (S2C Fig).

These findings suggest that *CLSTN2*, *DSCAM*, *GRIK1*, *CLOCK*, *INSL3*, and *NRXN3* are key candidate genes regulating male gonadal development. Collectively, these results indicate that the genes and biological functions regulating germ cell development differ across time points and between sexes. It is speculated that 55 dpc represents a critical period for sex determination and cell differentiation, while germ cells at 60 dpc are predominantly in the meiosis/mitosis transition stage

### Recapitulation of Goat Embryonic Gonadal Somatic Cell Development Based on Key Cell Trajectory Changes and Interactions

By examining the histological changes in gonads at different developmental stages and considering cellular heterogeneity, the differences and similarities between intersex, female and male goat embryos, as well as their underlying molecular mechanisms, were explored. Phenotype observations revealed that at 55 dpc, no significant morphological differences were observed between intersex and female fetuses, while male fetuses displayed the presence of a penis (S3A Fig). Histological analysis showed that XY male and XX sex-reversed gonads exhibited seminiferous cords differentiation and tunica albuginea formation. In contrast, XX female gonads exhibited the initial formation of the ovarian cortex surrounding a central undifferentiated embryonic structure (Fig 3B). These findings suggest that intersex gonads primarily exhibit a reduction in ovarian cortex and the formation and the presence of seminiferous cords and tunica albuginea, consistent with previous studies [46,47].

**Fig 3.**
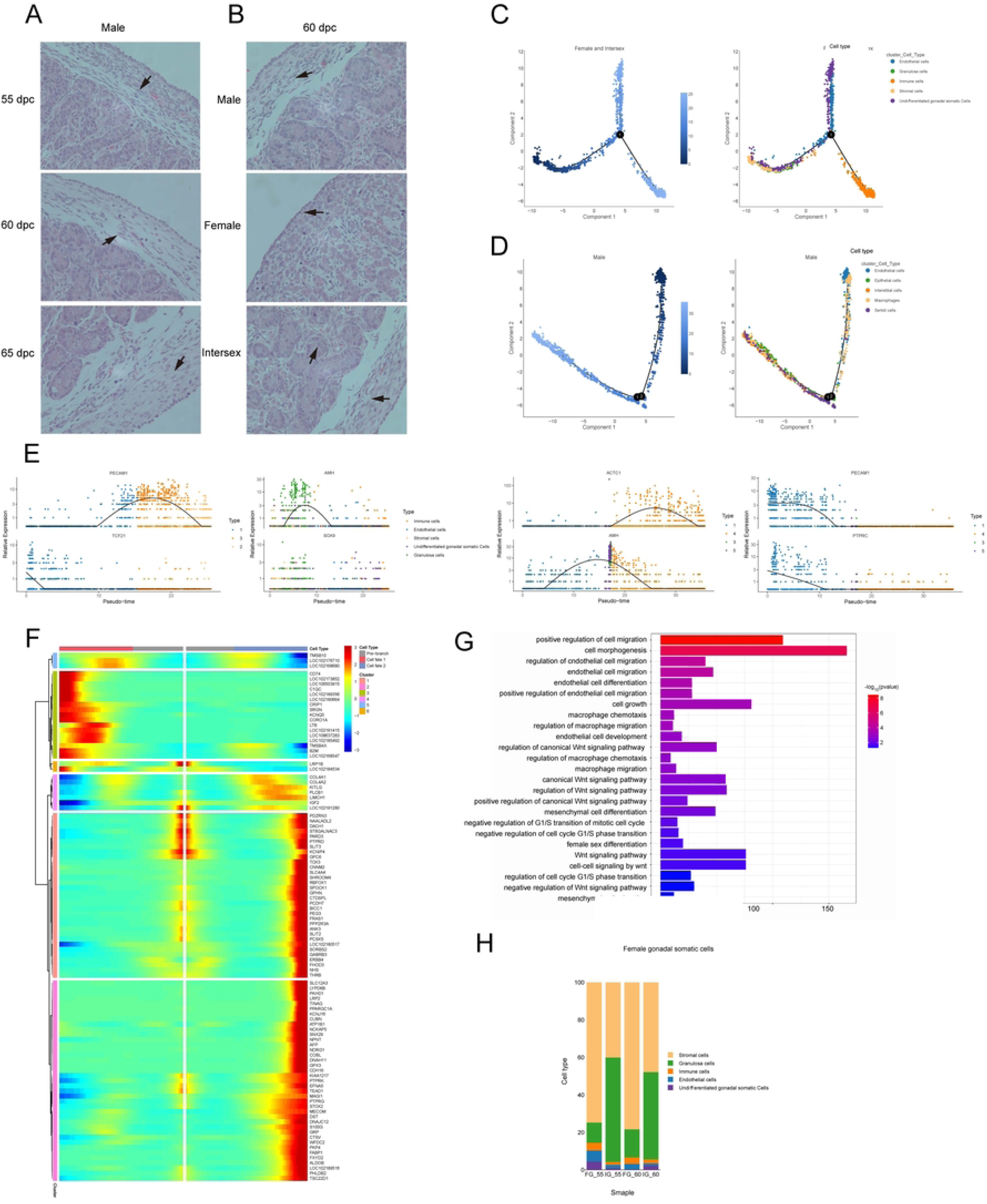
Recapitulation of development of goat embryonic gonadal somatic cells, and description of transcriptional features and developmental pathways in the fate determination of female and intersex gonadal somatic cells. **A**, Goandal tissue for 55, 60, and 65 dpc male fetuses. **B,** Gonadal tissue for female, intersex, and male at 60 dpc. **C-D,** Pseudotime trajectories (left) and developmental trajectories of different cell lineages along pseudo-time (right) of female, intersex (C), and male (D) gonadal somatic cells. Cell types were color-coded. **E,** The expression of maker gene was based on state and cell type. **F,** Clustering heat map showing the expression of branch node genes in female and intersex gonadal somatic cells. The top were branch nodes in the pseudo-chronological order, with each row representing a gene and each column representing a pseudo-time point, and the color represented the average expression value of the gene at the current point in time, ranging from red to blue, with decreasing expression levels. **G,** The main functions of GO enrichment differential genes of key branch in female and intersex gonadal somatic cells. **H,** Column diagram showing the difference in the proportion of female and intersex gonadal somatic cells. The colors corresponded to the cell types.

By 60 dpc, male fetuses displayed the presence of a scrotum and penis, and orchiocatabasis (testicular descent) was evident at 65 dpc. Compared with normal female fetuses, the 60 dpc intersex fetuses had a penis. The morphology of intersex fetuses remained similar to that of normal female fetuses (S3A Fig). Histological observations indicated that XY male gonads featured well-differentiated seminiferous cords and tunica albuginea, while XX female gonads featured well-developed ovarian cortex. In intersex gonads, the ovarian cortex was reduced, seminiferous cords were underdeveloped, and a thin layer resembling tunica albuginea separated the ovarian-like cortical regions (Fig 3A). These findings further confirmed that the reduction of the ovarian cortex and formation of seminiferous cords and tunica albuginea are key changes in intersex gonads [45,46].

To analyze key somatic cell types involved in gonadal development, single cell data were used to reconstruct pseudo-time differentiation trajectories via Monocle v.2.8.0 (S3C Fig and S3 Table). Analysis of dissociated single-cell subpopulations within the gonadal tissue revealed significant overlap across clusters, indicating their universal presence during gonadal development in dairy goats. Female and intersex embryos primarily exhibited one branch, three states, and five somatic cell types during early gonadal development (Fig 3C and S3B). In contrast male gonad cell lineages exhibited two branching points in the pseudo-time trajectory, uncovering five states and five somatic cell types (Fig 3D and S3B).

Pseudo-time trajectories for female and intersex gonads revealed that the stromal cell marker gene *TCF21* was exclusively expressed in state 1 during early gonadal development, suggesting stromal cells act as initial signaling hubs, relaying developmental signal to other key cells and stimulating granulosa cell migration and differentiation. The endothelial cell marker gene *PECAM1* exhibited high expression in the later part of state 1 as well as in states 2 and 3, indicating progressive proliferation and differentiation of endothelial cells during gonadal development (Fig 3E). The immune cell marker *AIF1* displayed dynamic expression patterns, particularly in state 2, suggesting a transient role in promoting gonadal development. Additionally, *SOX9* and *AMH* were significantly upregulated in state 1 and highly expressed in granulosa cells, indicating their pivotal role in the early formation of intersex gonads (Fig 3E).

In male gonadal trajectories, the macrophage marker genes *PTPRC* and *ACTC1* showed sustained high expression throughout development suggesting a critical role in early differentiation (Fig 3E). *PECAM1* and *KRT19* were highly expressed in state 1 (Fig 3E), indicating macrophage involvement in stimulating epithelial cell proliferation and differentiation. *AMH,* a marker for Sertoli cells, was highly expressed in states 3 and 5 (Fig 3E), whereas stromal cells were predominantly observed in state 4. The developmental trajectories of stromal and Sertoli cells exhibited substantial overlap, suggesting coordinated migration and differentiation processes in state 4, that facilitate male gonadal development.

In summary, the early embryonic gonadal development and the emergence of intersex characteristics are tightly linked to changes in cellular states and differentiation processes. The reduction of ovarian cortex formation and the emergence of seminiferous cords and tunica albuginea represent hallmarks of intersex gonadal development, with stromal, endothelial, and immune cells playing key roles in relaying developmental signals.

### Description of Transcriptional Features and Developmental Pathways in the Fate Determination of Granulosa Cells, Stromal Cells, and Endothelial Cells

To further investigate the developmental trajectories of key cell types, critical branches within the pseudo-time were analyzed. In the trajectories constructed for female and intersex goats, branch 1 consisted of six distinct gene sets, each representing three different developmental directions (Fig 3F and S4 Table). Analysis of these gene sets revealed significant differences in the expression levels of DEGs at various stages. High-expression genes from gene sets 3 and 5 were primarily found in cell fate 1, while those from gene set 6 were distributed across the pre-branch and cell fate 1. Genes from gene sets 1 and 4 were mainly concentrated in cell fates 1 and 2, whereas high-expression genes from gene set 2 were predominantly associated with cell fate 2 (Fig 3F).

t-SNE clustering analysis of female and intersexual gonadal cells, revealed that granulosa cells and stromal cells accounted for approximately 90% of somatic cells (Fig 3H). Further cell interaction analysis, using CellphoneDB software, showed that granulosa and stromal cells had stronger interactions with endothelial cells compared to other cell types (S3F Fig). This highlights the importance of characterizing the transcriptional features and developmental pathways of granulosa and stromal cells in relation to endothelial cells to understand the changes in intracellular expression dynamics during gonadal development.

GO enrichment analysis revealed significant enrichment in biological processes related to regulation of cell migration, endothelial cell differentiation, mesenchymal cell differentiation, and female sex differentiation (Fig 3G and S5 Table). Within gene set 2, genes such *LRP2* and *AFP* showed a marked upregulation during late gonadal development. In contrast gene set 1 was enriched for processes such as regulation of cell migration and female sex differentiation (S3H Fig). Both gene sets included key regulators of cell function, such as *DACH1*, *PHLDB2*, and *TEAD1*, which were highly expressed in endothelial, granulosa, and stromal cells. Dynamic expression patterns along the pseudo-time trajectory revealed the crucial regulatory roles of *TEAD1* and *PHLDB2* (S3H Fig). Therefore, genes including *LRP2*, *AFP*, *DACH1*, *PHLDB2*, and *TEAD1* are proposed as candidate regulators of granulosa, stromal, and endothelial cell fate determination.

Changes in gene expression are the primary drivers of the divergence of cell fates during gonadal development (S3G Fig). In cluster 2, genes such as *CTSS*, *TREM2*, and *PTPRC* were upregulated during the later stages of gonadal development. In cluster 3, *AFP* and *MEF2C* were predominantly upregulated in endothelial and immune cells during mid to late gonadal development (S3H Fig). Notably, the granulosa cell marker *AMH* and the endothelial cell marker *PECAM1* exhibited distinct dynamic expression changes, while the stromal cell marker *TCF21*, showed a downregulation during early gonadal development (Fig 3E).

Interestingly, *AMH* expression was significantly upregulated in granulosa cells of intersex embryos at 55 dpc, while *TCF21* was predominantly upregulated in stromal cells of female embryos. In contrast, *PECAM1* expression remained relatively stable in endothelial cells (S3F Fig). These findings indicate that the developmental pathways of granulosa, stromal and endothelial cells are dynamically regulated during goat gonadal development, with *AMH* and *TCF21* playing pivotal roles in the emergence of intersex characters.

### Delineating the Transcription Characteristics and Developmental pathways during Sertoli cell, Interstitial Cell and Epithelial cell Fate Decisions

In the male goat cell trajectory, both branches 1 and 2 comprised six distinct gene sets, each representing three different developmental directions (Fig 4A and S4A and S6 Table). In branch 1, highly expressed genes from gene sets 2 and 3 were primarily found in cell fate 1, while genes from gene sets 1 and 6 were concentrated in the pre-branch and cell fate 1. Highly expressed genes in gene set 5 were predominantly distributed in the pre-branch and cell fate 2, whereas gene set 4 genes were mainly located in cell fate 2. In branch 2, gene from gene sets 1, 2, and 5 showed high expression mainly in cell fate 2, while gene sets 4 and 6 were enriched in the pre-branch and cell fate 1. Genes from gene set 3 were specifically concentrated in cell fate 1.

**Fig 4.**
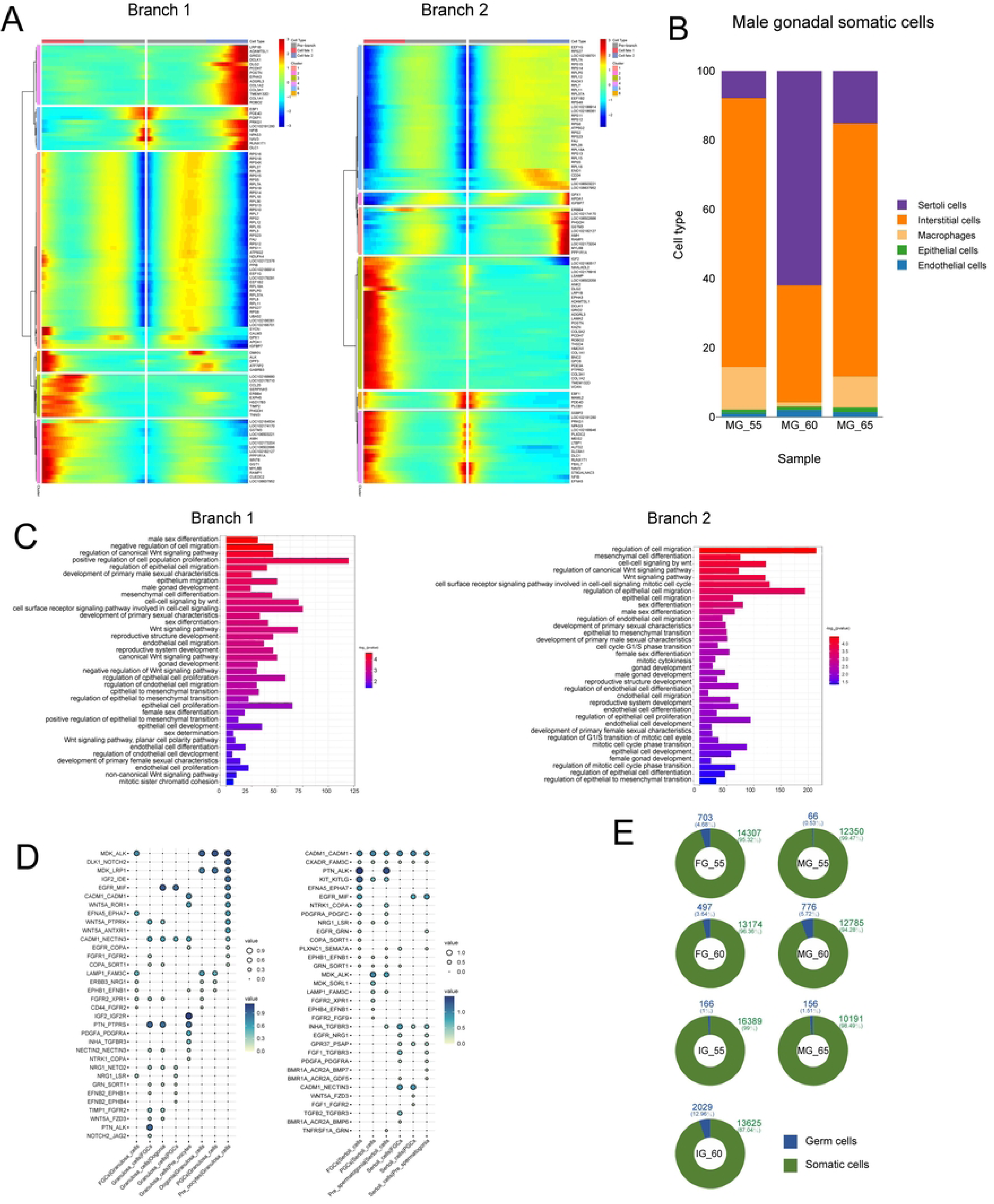
Delineating the transcription characteristics and development pathways during male gonadal somatic cell fate decisions, and revealing the characteristics of cell-cell interactions between gonadal somatic cell-germ cell. **A**, Clustering heat map showing the expression of branch node genes in male gonadal somatic cells. **B,** Column diagram showing the difference in the proportion of male gonadal somatic cells. The colors corresponded to the cell types. **C,** The main functions of GO enrichment differential genes of key branches in male gonadal somatic cells. **D,** Dot plot depicting representative ligand-receptor interactions between granulosa cells and germ cells (left), sertoli cells and germ cells (right). **E,** Doughnut diagrams showing the number and percentages of germ cells and somatic cells in each gonad. The colors corresponded to the cell types.

t-SNE clustering analysis revealed significant proportional differences in male gonadal cell types, although interstitial cells and Sertoli cells still constituted approximately 90% of all gonadal cells (Fig 4B). Cell interaction network analysis using CellPhoneDB demonstrated stronger interactions between Sertoli cells, interstitial cells, and endothelial cells compared to other cell types (S4B Fig). These findings prompted a focused investigation into the transcriptional and developmental pathways of these three lineages during gonadal differentiation.

To elucidate the developmental trajectories of Sertoli cells, interstitial cells, and epithelial cells during gonadal differentiation, GO enrichment and KEGG pathway analyses were performed on the DEGs from both branches were performed (S7 Table). Functional overlaps and differences were observed between branches 1 and 2. GO analysis revealed significant enrichment of biological processes, including regulation of cell migration, positive regulation of cell proliferation, male gonad development, mesenchymal cell differentiation, sex differentiation, and reproductive structure development (Fig 4C). KEGG analysis identified pathways such as the Wnt signaling pathway, cell cycle, estrogen signaling pathway, GnRH signaling pathway, circadian entrainment, circadian rhythm, and mitophagy-animal (S4C Fig). These pathways likely contribute to the differentiation of cells involved in gonadal formation and release of factors that stimulate cell migration and differentiation.

During gonadal development, processes such as gonadal differentiation, epithelial to interstitial cell conversion, and gonadal differentiation were significantly enriched. This suggests that some interstitial cells may originate from epithelial cells, influencing granulosa cell differentiation during gonadal development. The endothelial lineage also exhibited enrichment in relevant biological functions. The analysis further indicated that epithelial cells may transform into interstitial cells, thereby contributing to granulosa cell differentiation and influencing overall goat gonad development. Notably, *ARX* was exclusively expressed in interstitial cells (S4E Fig), and was significantly enriched in the regulation of epithelial cell proliferation within branch 2, indicating a close relationship between epithelial cells proliferation and interstitial cells development. Analysis of DEGs along the pseudo-time trajectory showed transient high expression of *SOX9* during mid-gonadal development, while *GATA4* exhibited consistent high expression during later developmental stages (S4D and S4F Fig). The expression trends of *SOX9*, *DMRT1*, and *GATA4* remained highly consistent across three male samples at 55 dpc, 60 dpc, and 65 dpc. Furthermore, genes such as *SOX4*, *SF1*, *NR5A1*, *TCF21*, *SMAD7*, *SNAI2*, *NOTCH1*, *WNT1*, and *WNT6* were enriched in GO terms linked to gonadal development (S4F Fig). Among these, *TCF21*, *SF1*, *ARX*, *NR5A1* and *SMAD7* were identified as potential candidate genes regulating male gonad development in goats.

### Revealing the Characteristics of Cell-Cell interactions Between granulosa Cell-Germ cell and Sertoli Cell-Germ Cell

Using CellPhoneDB, cell-cell interactions between granulosa cells, Sertoli cells, and germ cells were investigated to identify key ligand-receptor pairs involved in gonadal development. Several classical signaling pathways were conserved between germ cells and granulosa or Sertoli cells, including CXADR-FAM3C and CADM1-CADM1 (S8 Table). The expression patterns of receptors and ligands demonstrated cell type specificity. For example, IGF2-IDE and IGF2-IGF2R were specifically expressed between granulosa cells and pre-oocytes. Similarly, PTN-ALK interactions were observed between granulosa cells and FGCs, while ACVR-1B2B interactions were detected between granulosa cells and Oogonia.

Further analysis revealed additional signal interactions between PGCs, granulosa cells, and Sertoli cell. The SPP1-a4b1 complex and NRG1-LSR were identified as prominent interactions between PGCs and granulosa cells, while EFNA5-EPHA7 and TGFB2-TGFBR3 occurred between FGCs and Sertoli cells. Interactions such as MDK-SORL1 and WNT5A-FZD3 were detected between PGCs and Sertoli cells. Notably, *IGF2* and its receptor *IGF2R* exhibited enriched expression in granulosa cells and pre-oocytes, suggesting their pivotal role germ cells development.

In addition, compared with normal female fetuses with 55 dpc, the proportion of germ cells in intersex counterparts was lower (Fig 4E). Comparatively, the proportion of somatic cells in intersex counterparts was higher than that in normal female fetuses, indicating that there are some regulatory mechanisms in the gonads of fetuses that affects the development of somatic and germ cells, thereby regulating sex determination and gonad development.

### Two Testicle-Specific Macrophage Subpopulations identified in Fetal Goat Gonadal Cells

To investigate the role of macrophages in the testes, a subpopulation analysis based on the analysis of TREM2 expression defined two testis-specific macrophage populations: macrophage 1 (MA1) and macrophage 2 (MA2) (Fig 5A). In fetal goat gonads, 28.7% of macrophages were identified as MA1 (*TREM2^+^*), with the proportion of MA1 increasing with age (Fig 5B). Compared to MA2, the MA1 subpopulation exhibited greater heterogeneity, suggesting functional diversity within this group that the observed heterogeneity has functional implications for each cell (S5A Fig).

**Fig 5.**
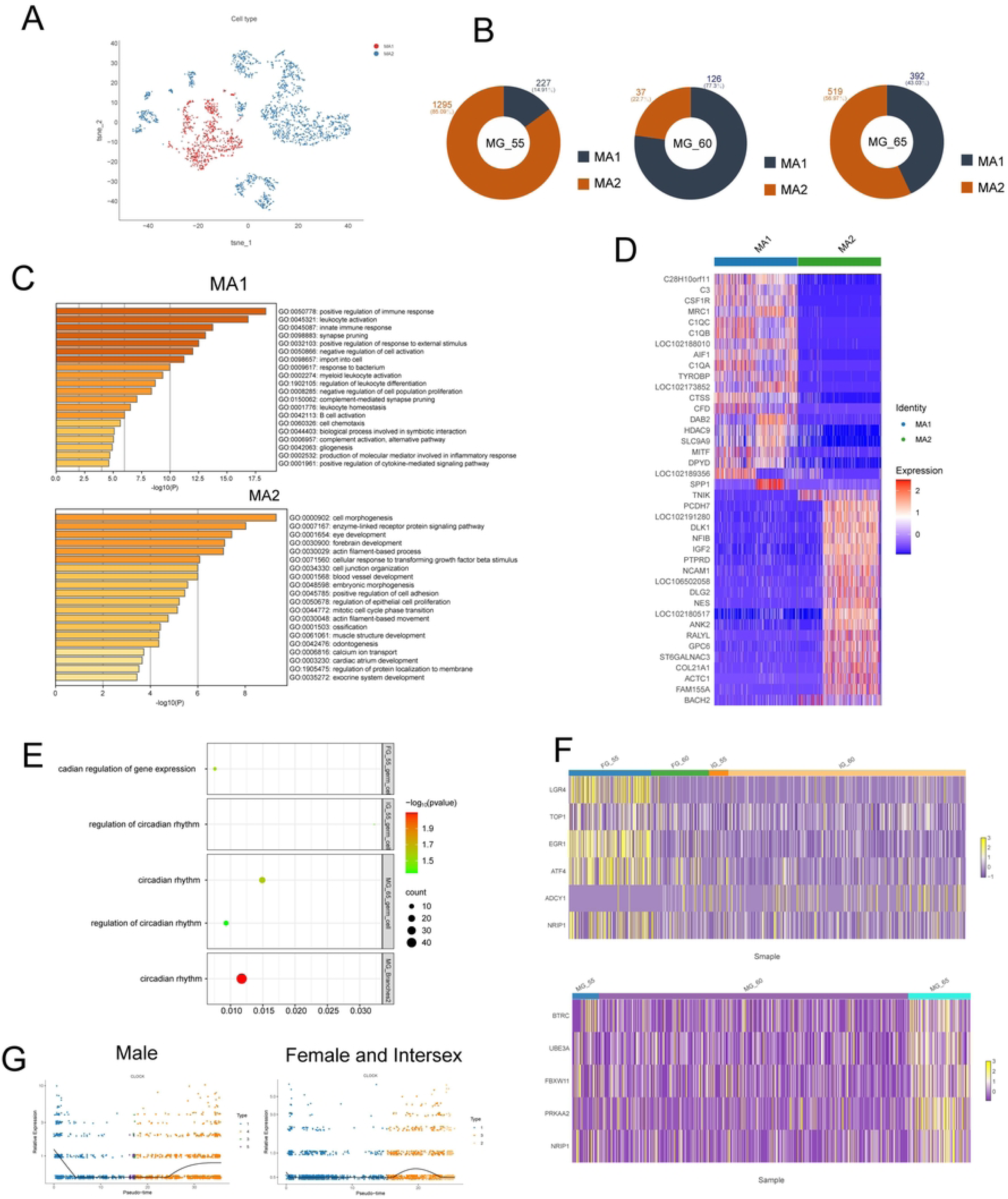
Two testicle-specific macrophages and circadian rhythm in the regulation of gonad development in goat embryos. **A**, t-SNE showing 2D visualization of macrophage subpopulations. Cells from cell subsets were color coded. **B,** Doughnut diagrams showing the number and percentages of MA1 and MA2 in the testis macrophages. The colors corresponded to the cell types. **C,** The main function of GO enrichment differential genes of MA1and MA2. **D,** T20 high expression genes in MA1 and MA2 cell subsets. **E,** Enriched GO function of gene related to circadian rhythm. **F,** Heat maps showing the expression of rhythm genes in female, intersex and male gonads. Each row represented a gene and each column represented a smaple. The color represented the average expression value of the gene at the smaple, ranging from yellow to purple, with decreasing expression levels. **G,** The expression of CLOCK gene was based on state.

To characterize the functional roles of these subpopulations, GO enrichment analysis was performed for the top 100 highly expressed genes (S9 Table). In MA1 subpopulation, enriched processes include positive regulation of immune response, leukocyte activation, innate immune response, synapse pruning, and positive regulation of response to external stimuli (Fig 5C and S5B and S10 Table). *TREM2* was notably enriched in these pathways, highlighting the role of the MA1 subpopulation in regulating immune and inflammatory responses. In contrast, the MA2 subpopulation was associated with cell morphogenesis, enzyme-linked receptor protein signaling pathways, actin filament-based processes, cellular responses to transforming growth factor beta stimuli, and cell junction organization (Fig 5C and S5B and S10 Table), with *COL1A2* being particularly enriched. Both subpopulations contributed to shared biological processes such as negative regulation of immune system processes, endocytosis, and positive regulation of *MAPK* cascades. Candidate genes for the MA1 subset included *C3*, *CSF1R*, *C1QC*, *C1QB*, *AIF1*, *C1QA*, and *TYROBP* (Fig 5D), suggesting that MA1 may contribute to maintaining the immune-privileged environment of the testes. In contrast, genes such as *PCDH7*, *NFIB*, *IGF2*, *PTPRD*, and others were highly expressed in the MA2 subset, indicating their involvement in cell surface signaling and cell morphogenesis.

### Circadian Rhythm May Regulate Gonadal Development in Goat Embryos

GO enrichment analysis of goat gonadal somatic and germ cells identified multiple terms related to circadian regulation of gene expression, regulation of circadian rhythm, and circadian rhythm (Fig 5E and S5C and S11 Table). Germ cells showed more significant enrichment in circadian regulation processes compared to somatic cells. Specifically, in MG_65 germ cells, *CLOCK* and *PER3* were enriched, while *PER1* and *PER2* were enriched in MG_55. In female gonadal somatic cells, genes such as *EGR1*, *LGR4*, *ATF4*, *TOP1*, and *NRIP1* were significantly upregulated in FG_55 (Fig 5F). Compared to normal female gonads, *ADCY1* was upregulated in IG_55. *PRKAA2*, *NRIP1*, *BTRC*, *FBXW11*, and *UBE3A* were upregulated in MG_65 and FGCs relative to normal female gonads (Fig 5F). These results suggested a significant role for circadian regulation in the development of goat gonadal germ cells across different sexes and stages.

In male gonadal somatic cells, several rhythmic genes including *CLOCK*, *PER2*, *PER1*, *CRY1*, and *CRY2*, were enriched, with *CLOCK* showing high expression during both early and late developmental stages (Fig 5G). Additionally, *ITPR1*, *GRIA1*, and *CACNA1D* were upregulated in male gonads at 65 dpc across different time points. For female and intersex gonadal somatic cells, branch 1 exhibited enrichment for *PER1*, which showed higher expression levels during early development (S5D Fig). *CLOCK* expression was elevated during later development, while *CRY1* was more highly expressed at 60 dpc compared to 55 dpc.

In intersexual gonads, *ARNTL* and *STAR* were upregulated, primarily in granulosa cells, compared to females. In contrast, *ATG5* and *CLOCK* showed higher expression in normal female gonads relative to intersexual ones (S5D Fig). These results indicated that *ARNTL* and *STAR* may be related to intersexual gonad development, while *ATG5* and *CLOCK* may promote normal female gonad development. Candidate genes enriched in the circadian entrainment pathway, including *GNAI2*, *GNG2*, *ADCY2*, *PRKG1* and *PLCB1* were up-regulated in females compared to intersex ones (Fig S5D). The expression of *PRKG1* and *PLCB1* displaying dynamic patterns during female and intersex gonadal development, while *GNAI2* and *GNG2* showed higher expression during the late development stage. *ADCY2* expression also increased at this stage (S5E Fig). These findings suggested a potential role of circadian regulation in the development and function of goat gonadal somatic cells.

### *FOXL2* and *SOX9* Participate in the Regulation of Gonad Development

To investigate the regulatory mechanisms of *FOXL2* and *SOX9* in female, male and intersexual gonads, immunofluorescence analysis was performed on fetal goat gonads to observe fluorescence localization and intensity of the three sex groups. Additionally, FOXL2 protein and SOX9 protein levels in the gonads of normal female, male and intersex fetal goat were compared.

The results showed that FOXL2 protein exhibited the highest expression in female gonads, followed by intersexual gonads, and the lowest levels were observed in males (Fig 6A). In contrast, SOX9 protein showed the highest expression in male gonads, followed by intersexual gonads, with the lowest expression detected in female gonads (Fig 6B).

**Fig 6.**
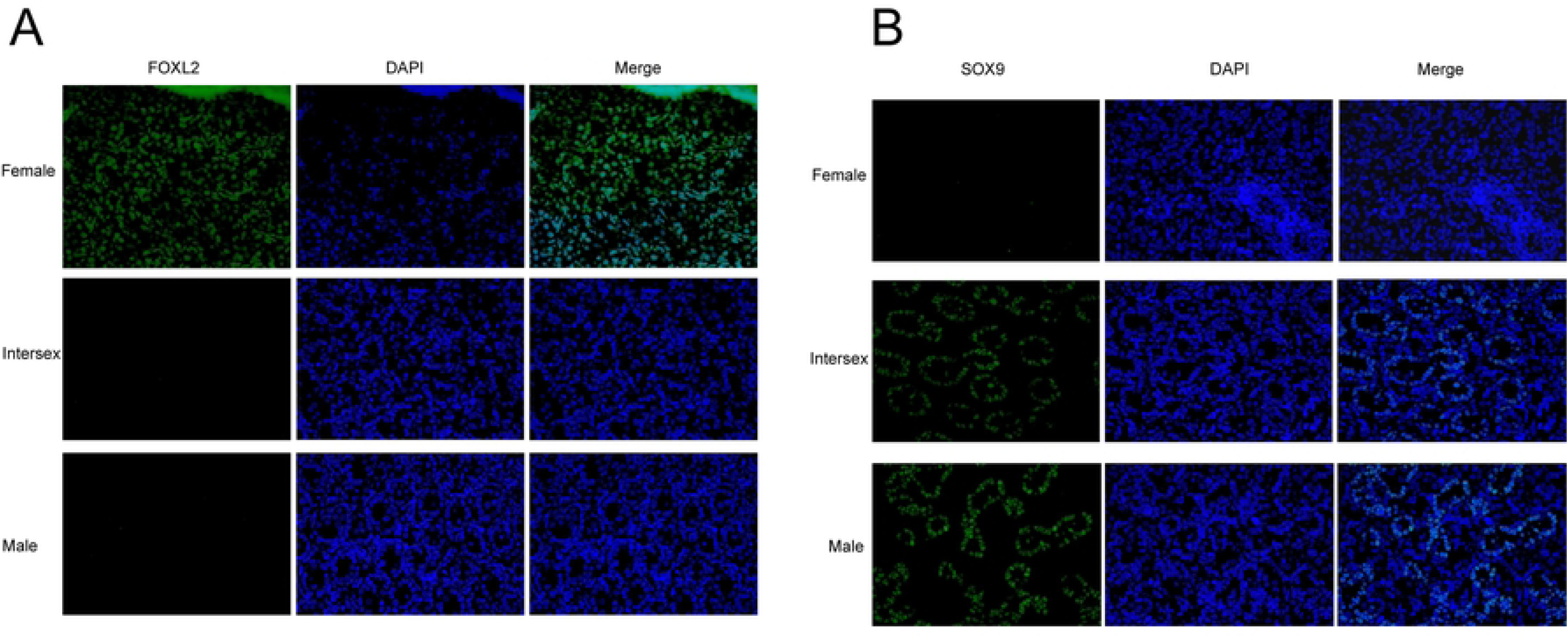
FOXL2 and SOX9 participate in the regulation of gonad development. **A-B**, Immunofluorescence analysis of FOXL2 (A) and SOX9 (B) in the gonads of goat.

Overall, the immunofluorescence results demonstrated clear differences in the protein expression levels of *FOXL2* and *SOX9* among female, male, and intersexual gonads, highlighting their contrasting roles in gonadal development and sex differentiation.

## DISCUSSION

### Revealing the Regulatory Mechanism of Intersex Formation in Goats Involving Different Cell Types

As an important livestock species, the regulatory mechanisms of germ and somatic cell development during sex determination in goats are not fully understood. In this study, we analyzed key gene expression profiles during goat embryonic gonad development, identified cellular heterogeneity, and constructed a single-cell map of early gonad development.

Our findings revealed that early goat gonadal cell types resemble those in other mammals, but differences appear in later stages. The origin of mammalian gonadal cells remains controversial, and may involve local mesenchymal cells, coelomic epithelial cells, or mesonephric cells [47–51]. Previous studies have shown that XX and XY embryonic gonads share lineage progression of supporting cells and the steroidogenic cells [52], which were differentiated into Sertoli and Leydig cells in the testis, or granulosa and theca cells in the ovaries [53]. Supporting-like cells (SLCs) were the first somatic lineage identified in bipotential gonads in mice at E10.5 and contributed to the pool of Sertoli and pre-granulosa cells [54].

Consistent with earlier findings, our study confirmed that *SOX9* can activate transcription of *FGF9* to initiate the differentiation of undifferentiated stromal cells into Sertoli cells [55]. In pseudo-temporal differentiation trajectories of female and intersex gonads, stromal cells appeared early and may act as initial signal for female gonad development. These cells subsequently stimulated the differentiation of granulosa cells, initiating gonad development and, in some cases, gender remodeling.

In male gonads, epithelial cells emerged as a potential source of gonadal cells, differentiating into interstitial cells. Additionally, macrophages played a key role in the early gonadal differentiation. Previous research indicated that coelomic epithelial cells were a source of testicular Sertoli and somatic cells in developing mouse gonads (11.2-11.4 dpc), although the ability was restricted at later stages [38,47,48]. However, XX goat gonads, showed no significant fate limitation for coelomic epithelial cells during the same developmental period [47,48].

According to this study, germ cells and somatic cells in the gonads were involved in both sex determination and gonad development. At 55 dpc, the proportion of germ cells detected in the ovaries was higher in the testes or XX sex-reversal gonads. This suggested that XX sex-reversed in XX (PIS^-^

^/-^) gonads occurred before the normal ovarian germ cells proliferative stage [35] and also before 55 dpc in fetal goats. Compared to normal ovaries, intersex gonads exhibited a higher proportion of granulosa cells and a lower proportion of, stromal cells. It was speculated that this imbalance may result from the differentiation of undifferentiated stromal cells into Sertoli cells [55], or the reprogramming of granulosa cells. These results suggested that the sex reversal of XX PIS^-/-^ gonads occured before the ovarian germ cell proliferation stage, approximately around 55 dpc [44,56].

For the first time, we comprehensively mapped the molecular landscape of normal male, normal female, and intersex gonads, revealing major cell fate decisions during gonad development.

### Revealing the Expression of Key Genes in Germ Cell Differentiation During Fetal Gonad Development

Germ cells were identified based on clusters expressing marker genes previously reported in humans, mice, or other mammals [38,57–60]. The pluripotency gene *POU5F1* exhibited high expression levels in mitotic PGC subsets, consistent with findings in humans and pigs, while *DDX4* was highly expressed in the mitotic stage of all male PGC [38]. *DAZL*, a germ cell specific and meiosis related gene, exhibited lat-stage expression, aligning with studies in mice [61,62]. Notably, *DDX4* was upregulated following the downregulation of *POU5F1* during oocyte development in mice [61], further supporting the evolutionary conservation of these stages.

GO enrichment results revealed that DEGs in germ cells were enriched in pathways such as stem cell differentiation, sex differentiation, mitochondrial translation and regulation of cell cycle process, cell surface receptor signaling pathway involved in cell-cell signaling, regulation of Wnt signaling pathway. These pathways are critical for germ cell differentiation, gonad development and intersex formation. Genes such as *HSF2* and *FOXO1* were linked to germ cell development, while *YY1* was associated with germ cell differentiation and gonadal evolution [63–66]. Members of the SOX family have been shown to contribute significantly to germ cell development [67]. In our investigation, *SOX15* displayed dynamic expression during mid-gonadal development, contributing significantly to germ cell development. Additionally, *ZEB2* was involved in cell signaling, and *FGF12* promoted follicular development [68–70], while its downregulation may be associated with the emergence of intersexual gonads. The *NR6A1*, which plays a role in the onset of ovulation [71], may also be implicated in intersex formation.

### Revealing the Expression of Key Genes in Somatic Cell Differentiation During Fetus Gonad Development

Cell fate changes during gonadal development revealed DEGs at key branch points, reflecting heterogeneity in gonadal somatic cell populations. We validated genes involved in gonad differentiation and sex determination across mammals, identifying candidates likely associated with intersex formation in goats.

The XX (PIS^-/-^) goats emerged due to a mutation in the PIS region [44,71], which likely affected downstream gene expression, leading to gonadal dysplasia and intersex formation. *FOXL2,* a key transcription factor for ovarian differentiation is regulated by the PIS fragments [72]. Its deletion triggers the upregulation of *SOX9* in XX PIS^-/-^ gonads, initiating male-specific differentiation pathways [44,73]. This was confirmed by genetic knock-out experiments [35] and also through immunofluorescence assays in our study. In mice, *SOX9* activates the transcription of *FGF9* and *AMH*, forming a positive feedback loop that enhances its own expression and stability, thereby initiating the differentiation of undifferentiated stromal cells into Sertoli cells [55,74]. Compared to normal female gonads of goat embryos at the same developmental stage, *SOX9*, *FGF9*, and *AMH* were upregulated in intersex counterparts, especially in granulosa cells, suggesting that these three genes may participate in the female-to-male sex reversal process. Furthermore, *DMRT1*, essential for male development and Sertoli cell maintenance, was found to be upregulated together with *SOX9* intersexual goats at in 55 dpc supporting its role in intersex formation [27,28,30,32]. *SOX9* induced mouse fibroblasts to differentiate into Sertoli cells under the action of *DMRT1* [22,23,75]. During female sex reversal, *SOX9* could bind to accessible genomic regions with the help of *DMRT1* [26]. Overexpression of *TCF21* in primary embryonic ovarian cells could cause abnormal expression of *AMH*, leading to the differentiation of Sertoli cells in females [74,76–78]. *AMH* has been confirmed in this study to be significantly elevated in intersex individuals at 55 dpc. However, in this study, despite an increase in the proportion of Sertoli cells and a decrease in stromal cells in intersex goats compared to normal female goats, *TCF21* expression was upregulated in females relative to intersex gonads. Thus, whether there was a differentiation of stromal cells into Sertoli cells during the female-to-male sex reversal process required further investigation.

Gonadal somatic cells were primarily enriched in pathways related to the regulation of cell migration, endothelial cell differentiation, sex differentiation, Wnt signaling pathway, estrogen signaling pathway, and GnRH signaling pathway (Fig 4C and 5C). The Wnt signaling pathway has been shown to play a crucial role in granulosa cell differentiation, sex determination, and gonadal development [79–83]. The estrogen signaling pathway promotes the proliferation of granulosa cells and induced the proliferation of epithelial and stromal cells, facilitating the development of female ovaries in bivalves [84–86]. The GnRH signaling pathway has also been identified as enriched in gonadal development-related pathways [87,88].

Several genes potentially involved in gonadal development were identified. The *LRP2* gene plays a role in maintaining male development [89]. *SF1* has been established as a nuclear receptor that regulates reproductive development and steroidogenesis [90], interacting with *PITX2* and participating in the Wnt pathway to enhance granulosa cell production in dairy goats [52]. Additionally, *ARX* and *LHX9* have been recognized as regulatory factors in Wnt signaling and gonadal development [52]. Other genes such as, *PHLDB2*, *ZNRF3*, and *TEAD1* may contribute to sex differentiation [91,92].

Promoting Wnt/β-catenin signaling can impair interstitial cell function and reduced *INSL3* secretion [80]. However, whether the upregulation of *INSL3* expression under intersex conditions affects granulosa cell development via the Wnt/β-catenin pathway warrants further investigation. *RSPO1*, identified as an female-specific somatic gene [35], is associated with masculinization of female gonads due to defects of *RSPO1* and *WNT4* in mice [83,92–94]. In this study, we found no difference in *RSPO1* expression among the gonads of the three sexes at early stages (55 dpc). However, at 60 dpc, *RSPO1* expression was the highest in the female ovaries, followed by intersex gonads, and the lowest in the male testes. *RSPO2*, a known target of *FOXL2* [35]. Therefore, it is speculated that *RSPO1* may also be one of the downstream target genes of *FOXL2*, following the downregulation of *FOXL2* expression in intersex.

### Effects of Testicle-Specific Macrophages on Fetal Gonad development

In goats, we identified *TREM2^+^* macrophages, similar to those observed in humans [40]. Enrichment analysis revealed distinct regulatory functions between macrophages subgroups. The MA1 subgroup primarily regulated immunoreaction, inflammatory response while the MA2 subgroup was involved in cell morphogenesis and cellular responses to transforming growth factor beta stimuli. In developing human testes, *TREM2^+^* fetal testicular macrophages with a microglia-like profile were identified as key regulators of the immunomodulatory environment prior to puberty [40]. These macrophages interact with Sertoli cells and spermatogonia to facilitate the clearance of apoptotic germ cells [40].

We identified additional genes potentially involved in the immune privilege mechanism of the testis, including *CSF1R*, *C1QC*, *C1QA*, *TYROBP, NFIX* and *IGF2*. *CSF1R* functions as a receptor, influencing *INHBA* to maintain macrophage populations in the testis [95] and is also involved in the self-renewal of mouse spermatogonia stem cells (SSC) [96]. *C1QC* alleviates inflammatory responses [97], while *C1QA* is associated with tissue macrophages and plays a critical role in antimicrobial responses [98]. *TYROBP* regulates immune cell migration and modulation [99] and *NFIX* is implicated spermatogonia meiosis in mice [100]. Additionally, *IGF2* promotes granulosa cells migration and differentiation [101].

Interestingly *TREM2^+^* immune cells were also identified in female and intersexual gonads, where they highly expressed *CSF1R*, *C1QC*, *C1QA*, and *TYROBP*. However, the expression of these genes was downregulated in intersex gonads compared to control female gonads. Among these genes, *CSF1R* has been shown to regulate granulosa cell responses to gonadotropins and promote early embryonic development [102,103]. These findings suggest that immune cell dysfunction might also play a critical role in the female-to-male sex reversal process in goats.

### Effects of Rhythm Genes on Fetal Gonad Development

Biological rhythms referred to the periodic changes in physiological and behavioral activities in organisms, regulated by rhythmic gene expression through the biological clock [104,105]. This study observed circadian rhythmic changes during early goat gonadal development, revealing how rhythm genes influence mammalian gonadal development.

GO enrichment analysis revealed significant enrichment in terms related to circadian regulation of gene expression. Several key DEGs associated with gonadal development were identified. The Bmal1 protein is expressed in interstitial cells of mouse testes, where it regulates testosterone production [106,107]. *BMAL1* (*ARNTL*) expression is positively correlated with *STAR,* a gene regulating steroid biosynthesis in human luteinizing granulosa cells [108], constituted with our findings. *NR1D1* participates in the autophagy process of granulosa cells [109,110], while *LGR4* is critical for granulosa cell survival and female development [111]. *ATF4* activates autophagy in ovarian granulosa cells [112]. Additionally, the expression and activity of *TOP1* in *C. elegans* are related to chromosome segregation, germline proliferation, and gonadal migration [113]. Inhibition of *ATG5* regulates granulosa cells autophagy [109,110], and *NRIP1* is involved in oogenesis [114,115]. In this study, *ATG5* and *NRIP1* were downregulated in intersex individuals, suggesting suppression of the female developmental pathway. Moreover, *BTRC* and *FBXW11* regulate the transition of male germ cells from mitosis to meiosis by degrading *DMRT1* [116]. In intersex individuals, *BTRC* and *FBXW11* expression was reduced, while *DMRT1* was slightly elevated, suggesting a role in intersex gonad formation through *DMRT1* regulation.

### Intersexual Goats May Be a Good Model for Human Reproductive Defects

The role of macrophages in promoting testicular function have been widely discussed [117,118], though there remains debate over whether different subpopulations of macrophages contribute to specific testicular function [119–121]. Yolk-sac-derived macrophages represent the earliest testicular macrophages, whereas fetal hematopoietic stem cells (HSCs) give rise to adult testicular macrophages, both of which play distinct roles in testicular morphogenesis [119].

In this study, immune cells were found to play an indispensable regulatory role in the differentiation and development of male and female sex glands in early goat embryos, with *TREM2^+^* macrophages potentially playing a critical role. In humans, *SIGLEC15+* and *TREM2+* fetal testicular macrophages signal to somatic cells outside and inside the developing testis cords, respectively [40]. This similarity suggests common mechanisms underlying embryonic gonadal development between goats and humans, making goats an excellent model for studying human reproductive disorders.

Although mice have long been used as models for human gonadal diseases, significant differences between mice and humans [35,122] hinder a complete understanding of the mechanisms underlying human gonadal development. Despite their importance for studying gonadal diseases and infertility [40], mice fail to replicate certain human-specific processes. Additionally obtaining human gonadal samples remains extremely challenging [40]. As an alternative, goats with intersex or normal gonads could provide a valuable model for exploring human sex determination mechanisms.

The gonads of goats and humans can synthesize estrogen during the gonadic switch, a feature absent in early ovarian differentiation in mice [35]. Additionally, *FOXL2* serves as a female sex-determining gene in humans and goats, whereas in mice, it maintained female somatic cells postnatally [35,73]. These findings highlight that goats are a more suitable model for studying human reproductive biology than mice.

In the analysis of somatic cell communication during gonadal development, endothelial cells exhibited strong interactions with Sertoli cells/granulosa cells, and stromal/interstitial cells. These results suggest that endothelial cells play a crucial role in the differentiation and development of gonads, particularly in male fetal gonads. There was significant overlap in the ligand-receptor pairs involved in key cell interactions during gonadal development in both humans and goats. For example, VEGFA-FLT1 interactions were observed between granulosa cells and endothelial cells, and PROS1-AXL interactions occurred between Sertoli cells and endothelial cells [38].

In humans, sPAX8s cells which mediate the formation of testicular and ovarian networks interact with endothelial cells [40]. Endothelial cells are also located near *SIGLEC15+* macrophages in the testes where they regulate the formation of testicular cords [46]. Additionally, the migration of renal endothelial cells has been identified as a transient but essential process for testicular cord formation [40]. These observations suggest that endothelial cells could serve as an entry point for studying human gonadal development mechanisms through goat models.

Core clock genes showed diurnal variation across various tissues, including differences between brain and gonads, though mice adapt differently to such changes compared to humans [123]. Circadian mechanisms play an important role in sex determination and differentiation. Disruption of circadian rhythm can cause human reproductive disorders such as polycystic ovary syndrome (PCOS) and/or premature ovarian insufficiency (POI) [124,125]. For instance, deletion of the Cacna2d3 gene, which encodes a circadian rhythm protein, leads to primordial follicles formation and reduced fertility in mice [126]. Additionally, maladjustment of circadian rhythms has been shown to affect interstitial cells,

resulting in reduced testosterone production [127]. In this study, circadian rhythm mechanisms were enriched in intersex, female, and male goat embryonic gonads. These findings further support the potential of goats as a model for human reproductive disorders.

### Materials and methods Experimental animals

Eleven pregnant ewes of Capra Hircus were obtained from the Sijichun dairy goat farm in Jimo District (Qingdao, China). Goats were raised according to dairy goat breeding standards and the experimental procedures were approved by the Experimental Animal Management Committee of Qingdao Agricultural University (DKY20231024, Annex 1). Dairy goats were anesthetized with lidocaine hydrochloride at 55 dpc, 60 dpc and 65 dpc (days post coitum, dpc), and 7, 6 and 2 fetuses were delivered by cesarean section. The corresponding gonadal tissue was obtained by surgery. The genetic sex and PIS genotype of 15 fetuses were detected by PCR [128]. For selected 7 embryos, one gonad tissue of each pair was used for scRNA-seq, and the other was used for HE staining and immunofluorescence analysis, denoted by “FG-55” (female gonads at 55 dpc), “FG-60“ (female gonads at 60 dpc), “IG-55” (intersex gonads at 55 dpc), “IG-60” (intersex gonads at 60 dpc), “MG-55” (male gonads at 55 dpc), “MG-60” (male gonads at 60 dpc) and “MG-65” (male gonads at 65 dpc), respectively (S12 Table).

### Dissociation of Gonad to a Single-cell Suspension

The gonadal tissue was removed, cleaned with PBS, cut into 2-4 mm tissue blocks, and put into enzymatic hydrolysate. The enzymolysis liquid system as follows: collagenase I (2 mg/mL) (C2674, Sigma), CaCl2 (10 mM) (G0070, Solarbio), DMEM/F12 medium (Catalog No. C11330500BT, Gibco, Beijing, China). The gonadal tissue was enzymolized at 37℃ for 20 min, and then trypsin (0.25%) was added to re-suspend cell precipitation. DMEM/F12 (4 mL) was added to terminate the enzymatic reaction. Filtering was performed using a 40 um filters to remove clumps and debris. The survival rate of cells in the sample was higher than 85%, which was convenient for the construction of the next single-cell library.

### Single-cell library construction and sequencing

Goat gonad single cell suspensions were obtained through the above steps, and scRNA-seq libraries were prepared according to manufacturer protocol of Chromium Next GEM Single Cell 3ʹ Reagent Kits v3.1 (Dual Index) for single cell barcoding and library. 16 000 cells for each sample, the Beads and partitioning Oil was loaded onto a Chromium Chip G. Resulting single-cell Gel Bead-in Emulsion (GEMs) were collected and linearly amplified. The swatch was conducted in a C1000 Touch Thermal cycler as 72℃ for 5 min, 98℃ for 30 s, cycled 12 x: 98℃ for 10 s, 63℃ for 30 s and 72℃ for 1 min. Emulsions were coalesced using the Recovery Agent and cleaned up using Dynabeads. The 10x Genomic Chromium barcode system was employed to construct a 10x barcoded cDNA library. Sequencing was conducted using an Illumina Hi Seq X10 sequencer, generating paired-end reads of 150 bp (PE150) for subsequent analysis.

### 10x Genomics scRNA-seq data processing

The Cell Ranger software pipeline (v5.0.0) developed by 10x Genomics was employed to demultiplex cellular barcodes, align reads to the genome and transcriptome utilizing the STAR algorithm. The unique molecular identifier (UMI) count matrix was subsequently analyzed using the R package Seurat (v3.1.1). To eliminate low quality cells and likely multiplet captures, the criteria was applied to filter out cells with gene numbers less than 200 and more than 10,000, UMI less than 1000. The low-quality cells were discarded, where >20% of the counts belonged to mitochondrial genes. Following the application of these quality control (QC) criteria, the single cells were retained for further analysis. Library size normalization was performed with Normalized Data function in Seurat to obtain the normalized count. Specifically, the global-scaling normalization approach adjusted the gene expression measurements for each cell based on total expression, multiplied by a scaling factor (10,000 by default), and the outcomes were log-transformed.

### Cluster analysis

Top variable genes across single cells were identified using the method [129] and FindVariableGenes function in Seurat [130]. In this project, the top 20 principal components exhibiting the highest explanatory variance from the principal component analysis (PCA) results were selected for dimensionality reduction through the RunPCA function in Seurat. The singleR software employs the existing purified cell types as references, calculates the correlation score between each cell and the reference, and combines the subpopulation maker gene for cell types annotation. Female and intersexual germ cells: PGCs marker gene *POU5F1* [38], FGCs marker gene *DDX4* [38,40.57,60], Oogonia marker gene *DAZL* [38,60], Pre-oocytes marker gene *NR2F2* [131]. Male germ cells: PGCs marker gene *POU5F1* [38]. FGCs marker gene *ZBTB16* [59] and pre-spermatogonia marker gene *EIF2S2* [58]. Female and intersexual somatic cells: stromal cell marker genes *TCF21*, *PDGFRB* [38,126], granulosa cell marker genes *AMH* and *SERPINE2* [132,133], endothelial cell marker gene *PECAM1* [38], immune cell marker genes *AIF1* and *PTPRC* [38,132], undifferentiated gonadal cell marker gene *CITED2* [134]. Male somatic cells: epithelial marker gene *KRT19* [60], Sertoli marker gene *AMH* [38,135], macrophage marker genes *ACTC1* and *PTPRC* [136], and stromal cell marker genes *TCF21* and *PDGFRB* [38,136], endothelial cell marker gene *PECAM1* [38]. Marker genes were manually annotated by consulting the ZFIN database (http://zfin.org/) and Pubmed (https://pubmed.ncbi.nlm.nih.gov/) indexed literature on mammalian gonad development. Cellular visualization was used a 2-dimensional Uniform Manifold Approximation and Projection (t-SNE) algorithm with the RunUMAP function in Seurat.

### Pseudo-time trajectory analysis

Single-cell trajectories reconstruction analysis was performed using the Monocle 2 package (v2.8.0) for the discovery of cell state transitions. The following parameters were used: average expression R0.125, num_cells_expressed R10, and q_val<0.01. A branch-specific gene expression heatmap was plotted with plot_pseudotime_heatmap function.

### Enrichment Analysis

Differentially expressed genes (DEGs) were identified using the FindMarkers function [130]. A threshold of P-value < 0.01 and |log2foldchange| > 0.58 was set for significant differential expression. Gene Ontology (GO) and Kyoto Encyclopedia of Genes and Genomes (KEGG) were utilized for the biological process enrichment analysis in order to gain a deeper understanding of the functions of DEGs between Capra Hircus gonadal tissue. Enrichment analysis was conducted by using the “clusterProfiler” R package and Metascape. All analyses were considered to be significantly enriched if the p-value<0.05. The P-value was adjusted using Benjamini-Hochberg correction.

### Data and code availability

The dairy goat scRNA-seq data used in this study have been deposited in the Genome Sequence Archive in National Genomics Data Center, China National Center for Bioinformation/Beijing Institute of Genomics, Chinese Academy of Sciences (GSA: CRA020240) that are publicly accessible at https://ngdc.cncb.ac.cn/databasecommons/database/id/1790. All original code has been deposited at github and is publicly available at https://github.com/xin189/original-code.git.

### Cell interaction analysis and cell heterogeneity difference analysis

CellphoneDB [137,138] was used to analyze the communication networks between different cell types, including a database of ligands, receptors and their interactions. The Euclidean distance between all pairs of cells in the subpopulation was calculated and its distribution was expressed in the form of boxplot. The higher the distance distribution, the higher the cell heterogeneity of the subpopulation was [139].

### Hematoxylin-Eosin (HE) staining and immunofluorescence

Fresh gonads were collected and fixed in Bouin’s fluid for 24 h, flushed for 3 h. The fixed gonads were dehydrated, immersed for 15 min with xylene, entrapped in paraffin, and sectioned at 5 um. The section were stained with hematoxylin for 5 min, eosin for 3 min, and finally sealed with neutral balsam. Immunofluorescence analysis was conducted on the gonads of the three genders under identical conditions. Sections were baked in a constant temperature oven at 56℃ for 2 h, dewaxed and rehydrated. Then, sections were immersed in Citrate Antigen Retrieval Solution at 96℃ for 10 min, cooled at room temperature, and washed three times in PBS for 5 min each time. After 3-5% BSA incubation, sections were added primary antibody diluted with 3-5% BSA, and incubated overnight in a 4°C refrigerator. After incubation with primary antibody, sections were rewarmed at room temperature for 30 min, washed three times in PBS for 10, 20 and 20 min in sequence. Sections were treated with a secondary antibody, incubated at room temperature for 30 min, and washed in PBS three times for 5 min each time. Then DAPI was added, incubated at room temperature for 3 min, and with Antifade Mounting Medium were added, stored at 4℃. The pictures were captured using a Nikon AR1 confocal system (Nikon, Tokyo, Japan). Images were then taken with an Olympus BX51 microscope imaging system (Olympus, Tokyo, Japan).

## Supporting information

**S1 Fig. scRNA-seq delineated cellular heterogeneity during dairy goat gonad development. A,** t-SNE embedding visualization of germ cell clusters and gonadal somatic cell clusters from female, intersex, and male gonads based on scRNA-seq data. Cells from different clusters were color coded. **B-C,** Doughnut diagrams showing the number and percentages of transcriptionally identified cell populations in each of the gonadal germ cell. The colors corresponded to the cell types. **D,** t-SNE embedding visualization of gonadal somatic cells from macaque, pig, and goat gonads based on scRNA-seq data. Cells were color-coded according to standardized gene activity.

**S2 Fig. Dynamics gene expression and transcriptional regulatory network in germ cells of different ages. A,** Pseudotime trajectories of female, intersex (left), and male (right) germ cells. **B,** Heatmap showing the gene expression levels along the pseudotime trajectories in female and intersex germ cells. **C,** The expression levels of representative genes in the identified cell types were shown in male germ cells.

**S3 Fig. Recapitulation of development of goat embryonic gonadal somatic cells, and description of transcriptional features and developmental pathways in the fate determination of female and intersex gonadal somatic cells. A,** Apparent comparison of 55, 60, 65 dpc fetal goat. **B,** Clusters (left) and states (right) along pseudotime trajectories of female, intersex, and male gonadal somatic cells. Clusters and states were color-coded. **C,** Tree trajectory based on cluster. **D,** The expression levels of representative genes in the identified cell types were shown in male gonadal somatic cells. **E,** The expression levels of representative genes were shown in female and intersex gonadal somatic cells. **F,** Cell-cell interactions in female and intersex gonadal somatic cells. **G,** Heat map showing changes of branch node genes’ expression in female and intersex gonadal somatic cells. **H,** Visualization of expression levels of related genes that lead to different cell fates in female and intersex fetuses was based on state (left) and cell type (right).

**S4 Fig. Delineating the transcription characteristics and development pathways during male gonadal somatic cell fate decisions, and revealing the characteristics of cell-cell interactions between gonadal somatic cell-germ cell. A,** Heat map showing changes of branch node genes’ expression in male gonadal somatic cells. **B,** Cell-cell interactions in male gonadal somatic cells. **C,** KEGG analysis of male gonadal somatic cells. **D,** The expression levels of representative genes were shown in male gonadal somatic cells. **E,** The expression levels of *ARX* in the identified cell types were shown in male gonadal somatic cells. **F,** Based on cell type and state, visualization of expression levels of related genes that lead to different cell fates in male fetuses.

**S5 Fig. Two testicle-specific macrophages and circadian rhythm in the regulation of gonad development in goat embryos. A,** Heterogeneity of MA1 and MA2 cell subsets. The higher the distance distribution, the higher the cell heterogeneity of the cell subset. **B,** Network layout showing representative terms of gene function enrichment in MA1 and MA2. Circle node represented each term, where its size was proportional to the number of input genes fall under that term, and its color represented its cluster identity. **C,** KEGG analysis of gene related to circadian rhythm. **D,** The expression levels of circadian rhythm related genes at different time points in female and intersex gonads. **E,** Visualization of expression levels of circadian rhythm genes regulating gonad development in female and intersex fetuses.

**S1 Table. Number of gonadal germ cells and somatic cells after stringent quality control in different sexes and different ages.**

**(XLSX)**

**S2 Table. Enrichment of gonadal germ cells’ DEGs and their associated biological processes at various ages and across sexes.**

**(XLSX)**

**S3 Table. Genes involved in constructing pseudo-time differentiation trajectories.**

**(XLSX)**

**S4 Table. Six distinct gene sets in pseudo-time differentiation trajectories constructed by female and intersexual goat gonads.**

**(XLSX)**

**S5 Table. Enrichment of gonadal somatic cells’ DEGs in branch1 of pseudo-time differentiation trajectories constructed by female and intersexual goats.**

**(XLSX)**

**S6 Table. Six distinct gene sets in pseudo-time differentiation trajectories constructed by male goat gonads.**

**(XLSX)**

**S7 Table. GO enrichment and KEGG pathway of gonadal somatic cells’ DEGs in pseudo-time differentiation trajectories constructed by male goats.**

**(XLSX)**

**S8 Table. Key ligand-receptor pairs involved in gonadal development.**

**(XLSX)**

**S9 Table. DEGs for identified Macrophage Subpopulations.**

**(XLSX)**

**S10 Table. Gene Set Enrichment Analysis of identified Macrophage subpopulations.**

**(XLSX)**

**S11 Table. GO enrichment and KEGG analysis of goat gonadal somatic and germ cells revealed terms related to circadian rhythm.**

**(XLSX)**

**S12 Table. Basic information of goat embryos.**

**(XLSX)**

**S1 Appendix. The Animal Ethics Procedures and Guidelines of the People’s Republic of China.**

**(PDF)**

## Author Contributions

**Conceptualization:** Hegang Li, Jinshan Zhao **Formal analysis:** Hegang Li, Jinshan Zhao **Funding acquisition:** Jinshan Zhao, Hegang Li **Investigation:** Hegang Li, Jinshan Zhao **Methodology:** Hegang Li, Xinxin Cao, Yichen Zhang, Dongliang Xu, Mengmeng Du, Xiaokun Lin, Lu Leng, FM Perez Campo, Lihui Zhang, Mingzhi Sun, Xiaoxiao Gao, Jianning He, Qinan Zhao, Jianguang Wang **Project administration:**Hegang Li, Jinshan Zhao **Supervision:** Hegang Li, Jinshan Zhao **Validation:** Hegang Li, Jinshan Zhao **Visualization:** Xinxin Cao, Mengmeng Du **Writing – original draft:** Xinxin Cao, Hegang Li, Yichen Zhang **Writing – review & editing:** Hegang Li, Jinshan Zhao, FM Perez Campo

